# Cryo-EM reveals the central steps of mitochondrial complex III assembly and the cooperative assembly of supercomplex CIII_2_CIV

**DOI:** 10.64898/2026.09.07.749710

**Authors:** Ahad Ali Kazmi, Andrea Gottinger, Marine S. Huhardeaux, Andrea Stärk, Beatrix Santiago-Schübel, Emma J. Fenech, Irene Vercellino

## Abstract

Mitochondrial complex III is the central component of the respiratory chain and is conserved across eukaryotes. Complex III is an obligate dimer which assembles through a stepwise mechanism, involving 20 subunits and other assembly factors. Defects in the assembly of complex III are associated with metabolic diseases. The assembly mechanism of complex III has long been investigated using biochemical approaches, which suggested a process where folded subunits are added sequentially and in parallel for the two protomers after dimerization. Our structural investigation of CIII_2_ assembly challenges these assumptions: we observe using cryo-EM that incorporation and folding of its subunits can be uncoupled (as we show for cytochrome c_1_) and that after dimerization the assembly of the complex does not proceed in parallel for the two protomers (as we show for the folding of the intermembrane space domain). Our structures also reveal the mechanism of formation of supercomplex CIII_2_CIV for non-vertebrates, intertwined with the assembly of CIII_2_. This work reshapes our knowledge of complex III assembly and proposes a generalizable model for the maturation of the complex, alongside a model for supercomplex formation that supports the cooperative assembly model. Furthermore, as the observed assembly steps cannot be predicted by AlphaFold, our work also showcases the central role of cryo-EM in the study of assembly mechanism of protein complexes.

## Introduction

Oxidative phosphorylation (OXPHOS) is a key process in aerobic respiration that produces adenosine triphosphate (ATP) to fuel cellular chemical reactions and sustain life. OXPHOS produces an electrochemical proton gradient across membranes to drive ATP synthesis via ATP synthase (complex V) by harvesting the free energy from the transfer of electrons through the respiratory chain^1^. The respiratory chain is an enzymatic system that is extremely conserved across the tree of life. In eukaryotes, respiratory chain enzymes, or complexes, reside in the inner mitochondrial membrane (IMM). Canonical eukaryotic respiratory chains are composed of four respiratory complexes (CI, CII, CIII, CIV) and two mobile electron carriers, IMM-bound quinone and soluble cytochrome c (Cyc1) in the intermembrane space (IMS). CI and CII, alongside additional enzymes, reduce quinone to quinol by transferring electrons from NADH and succinate respectively. Upon quinol binding, CIII oxidizes it to quinone while reducing Cyc1. CIV transfers electrons from reduced Cyc1 to the terminal electron carrier, molecular oxygen, thereby completing the chain^2^. CIII hence occupies a central position in the respiratory chain by bridging the flow of electrons from reducing equivalents formed in the enzymatic reactions of CI and CII (NADH and succinate) to molecular oxygen that binds CIV.

CIII is only catalytically active once it is fully assembled as an obligate homodimer composed of 10 subunits per protomer, therefore it is commonly referred to as CIII_2_^3^. Despite the central role of CIII_2_ in the respiratory chain, its extreme evolutionary conservation and the occurrence of severe metabolic diseases^4–6^ when its assembly is perturbed, the mechanism of CIII_2_ formation is still incompletely resolved, in particular regarding the intermediate steps, for which no consensus on the order of events has been reached and no assembly factors have so far been identified^7^. Our current knowledge of CIII_2_ assembly derives primarily from biochemical studies of mammalian and yeast systems, where CIII_2_ assembly was stalled to accumulate intermediates. CIII_2_ assembly was proposed to be assisted by assembly factors in the initial and final stages and to start in a stepwise manner with the synthesis of cytochrome b (Cob), the subsequent addition of Qcr7 and Qcr8, the maturation through insertion of Cytochrome c_1_ (Cyt1), Qcr6, the Core 1 and 2 (Cor1 and Cor2) proteins, alongside the dimerization of the complex and the final incorporation of the catalytic subunit iron sulphur protein (ISP, hereby referred to as the Rieske protein, Rip1), together with Qcr9 and Qcr10^8–19^. These studies assumed that CIII_2_ subunits are incorporated as fully folded proteins in the nascent complex and were not able to conclusively elucidate the intermediate steps of CIII_2_ assembly, between insertion of Qcr7 and the final addition of Rip1, Qcr9 and Qcr10.

Furthermore, although the respiratory complexes are functional in isolation, it is well established that in the mitochondrial membrane they associate into higher-order structures called supercomplexes^2^. In *S cerevisiae*, supercomplexes CIII_2_CIV_2_ and CIII_2_CIV have been identified, where the obligate complex III dimer is flanked by either one or two copies of complex IV^20,21^. These supercomplexes assemble exclusively with the canonical subunits of CIII_2_ and CIV, without requiring additional specific assembly factors. Conversely, in vertebrates the formation of CIII_2_CIV is mediated by the assembly factor SCAF1, which then becomes a *bona fide* subunit of the supercomplex, in addition to mediating the formation of a CICIII_2_CIV_2_ supercomplex named the SC-respirasome^22,23^. The assembly mechanism of supercomplexes has long been a subject of debate, centered on two alternative models: one where the single complexes have to first fully assemble before forming supercomplexes and one where assembly intermediates of the complexes already combine into supercomplexes^2^. Structural biology and biochemistry, primarily from mammalian systems favor the latter model, leading to the question whether the cooperative assembly is a common feature of the respiratory chain in eukaryotes.

Here, by using cryo-electron microscopy, we present the structures of CIII_2_ and CIII_2_CIV assembly intermediates from a *Saccharomyces cerevisiae (S. cerevisiae)* strain where CIII_2_ assembly is stalled since the late assembly catalytic subunit Rip1 is knocked-out (Rip1KO). Our cryo-EM data of the CIII_2_ assembly intermediates propose a conserved assembly mechanism and challenge the notion that fully folded CIII_2_ subunits are incorporated in the nascent complex. Instead, the structures indicate a more intricate scenario where IMS-facing subunits Qcr8 and Cyt1 partially fold in their transmembrane regions, thereby stabilizing CIII_2_, followed by the folding of the soluble domains of both proteins, and the concomitant incorporation of Qcr6. Thus, the folding of the transmembrane regions of the IMS-facing subunits is required for CIII_2_ stability during assembly, whereas the folding of the soluble domains of the same proteins is not. Structural analysis of the CIII_2_CIV intermediates revealed that a fully assembled complex IV binds to a CIII_2_ intermediate that is at a later assembly stage than the CIII_2_ species observed in isolation, where Qcr8 and Cyt1 are fully assembled. Based on our structures, complex IV first associates with CIII_2_ and then changes conformation towards the position of the fully assembled supercomplex. These results are the first description of the assembly of supercomplex CIII_2_CIV in non-vertebrates, where the supercomplex assembly factor SCAF1 that mediates the supercomplex formation in vertebrates^22^ is absent. Despite the divergent mechanism of CIV association with CIII_2_, which depends on the presence or absence of SCAF1, our structures and the mammalian structures converge on supporting the cooperative assembly model of supercomplexes and also converge on the conserved assembly mechanism of CIII_2_, featuring the same assembly intermediate of CIII_2_ bound to CIV before the last step of CIII_2_ maturation.

## Results and discussion

### The Rip1KO forms a stable and native-like CIII_2_ intermediate

To elucidate the assembly mechanism of CIII_2_ we made use of a published strain, where Rip1 is KO^10^, which allows the accumulation of assembly intermediates at steady state. Although the Rip1KO grew slower than the wild-type (WT) and plateaued at lower optical density (OD), the growth curve still displayed the logarithmic phase similarly to the WT counterpart, providing enough material at early logarithmic phase for structural analysis (Fig 1G). Our cryo-EM data analysis of the purified CIII_2_ revealed structures of CIII_2_ in supercomplex (SC) with CIV (Fig 1A-1C) and in isolation (Fig 1B-1D). In both cases, the consensus maps expectedly showed no density for Rip1, alongside no density for Qcr10 and partial density for Qcr9. This intermediate perfectly recapitulates the intermediate observed from wild type mice^22^, suggesting that the disruption of Rip1 in the mutant strain does not affect the formation of a native-like CIII_2_.

**Figure 1.**
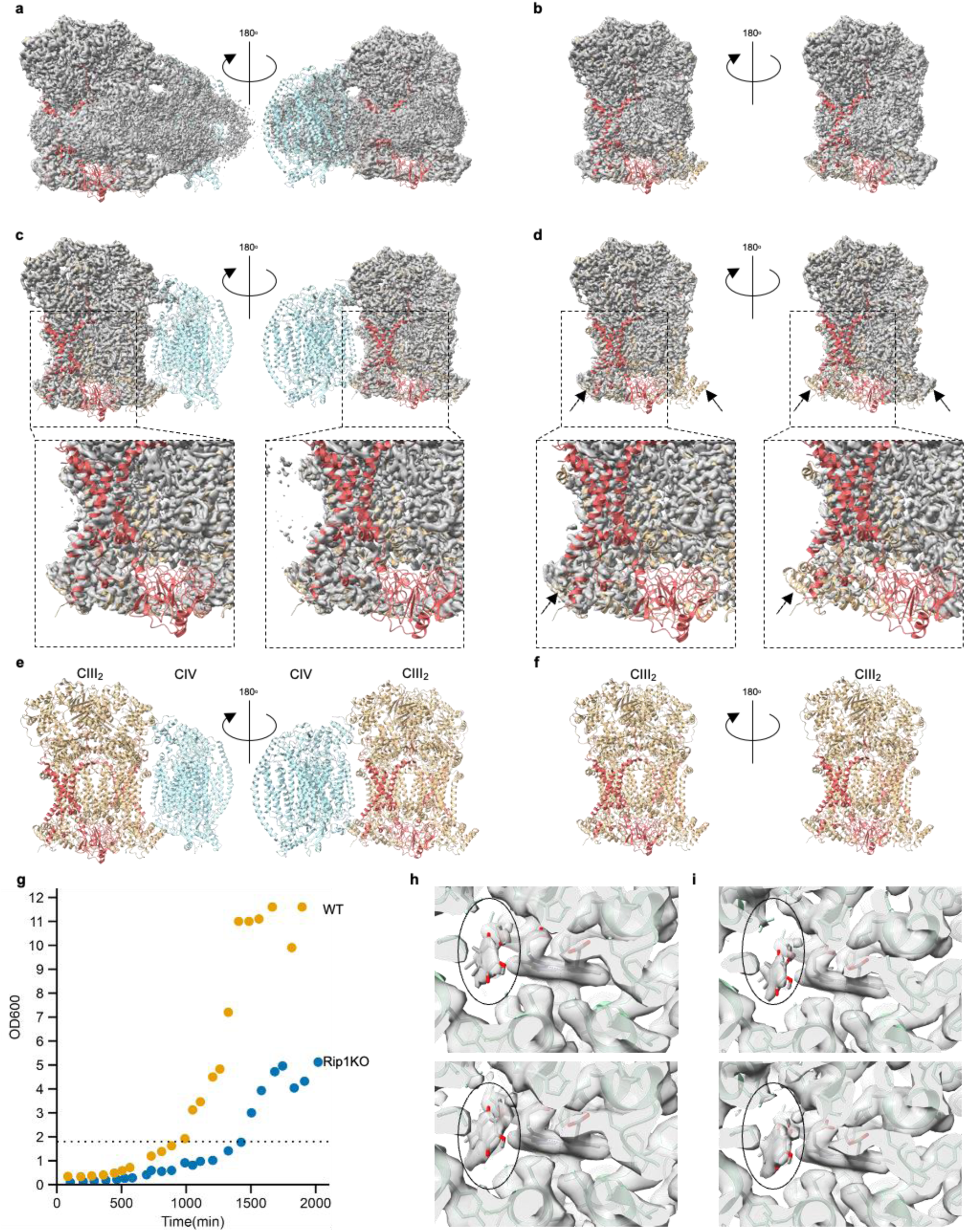
CIII_2_ from Rip1KO strain retains a native-like structure in isolation and in supercomplex. a-d. Consensus maps of supercomplex CIII_2_CIV (a and c) and isolated CIII_2_ (b and d) at low (a and b, map level 0.05) and high (c and d, map level 0.08) stringency, featuring fitted models for CIII_2_ (tan and Indian red) and CIV (powder blue). The arrows in d highlight density differences in the Qcr6 area. **e-f**. Fitted CIII_2_CIV and CIII_2_ models without maps. The model is the fully assembled CIII_2_CIV (pdb 9etz), with CIV in blue, CIII_2_ in tan except for subunits Qcr9, Qcr10 and Rip1 which are in red. **g.** Growth curve of WT and Rip1KO yeast. **h-i.** Cob (coloured green in the model), heme b_H_ and quinone (both coloured grey by atom, quinone encircled) density in the Qi site in the two protomers of isolated CIII_2_ (h) and CIII_2_ from SC (i).

Furthermore, although the intermediate is catalytically inactive due to the absence of the electron-transferring subunit Rip1, we observed clear density for the head group of quinone, the electron acceptor of the low potential heme b_H_, in the Q_i_ site. Although quinone is not constantly bound to CIII_2_ since it is released in the membrane after its reduction during the catalytic cycle of the enzyme (Q Cycle)^3,24–28^, it is featured bound to the Qi site in several X-ray and cryo-EM structures ^20,21,29–34^. Our data now reveal that this relatively stable interaction of quinone with Cob in the Q_i_ site is not exclusively a feature of the fully assembled, catalytically competent CIII_2_, but it is already present during assembly when the complex per se is inactive, harbors a fully folded quinone binding site. This feature might suggest that CIII_2_ is assembled in a catalytically ready state and once fully assembled, it can immediately start the Q cycle upon quinol binding in the Q_o_ site.

When assessing the map features, we observed that the CIV density in the supercomplex is weaker than CIII_2_, indicating conformational variability (Fig 1A-1C), while the isolated CIII_2_ intermediate shows better density for Qcr6 in one protomer than in the other (Fig 1B-D and arrows), pointing to potential compositional variability in the CIII_2_ structures. We therefore set to further classify the CIII_2_ and supercomplex particles to separate the different species present in each sample.

### The IMS domain of CIII_2_ assembles one protomer at a time

Thorough classification of the isolated CIII_2_ particles separated four different intermediate states representing the folding of the intermembrane space (IMS) domain of CIII_2_ (Figs 2-3). As shown in Fig 2A (surface representation, coloured tan), all the intermediates feature a core CIII_2_ dimer comprising the soluble subunits Cor1 and Cor2 on the matrix side and the membrane-bound subunits Cob, Qcr7, Qcr8 in an almost perfectly dimeric form, alongside the transmembrane helix of subunit Cyt1 (Fig 3A). This core structure represents the earliest intermediate, which we name intermediate I and is followed by three subsequent intermediates where the IMS domain assembles asymmetrically around the soluble domain of Cyt1. Specifically, in Intermediate II the soluble domain of one Cyt1 protomer is ordered and encased by the neighbouring Qcr6, the previously disordered C-terminus of Qcr8 and the previously unresolved Qcr9, now folded in the C-terminal region (Figs 2-3). In the subsequent Intermediate III, the Cyt1 soluble domain from the second protomer is mainly ordered and flanked by the second copy of Qcr9, visible in the C-terminal region. The appearence of the density for the second Cyt1 soluble domain is also accompanied by ordering of the 135-146 loop of Cyt1 from the first protomer (full circle in Fig 3A), while the N-terminus (until residue 75) and the 198-212 loop of Cyt1 from the second protomer remain unresolved in Intermediate III and are only stabilized by the interaction with the second copy of Qcr6 (dotted circle in Fig 3A) in Intermediate IV, which also coincides with folding of the C-terminus of the second Qcr8 copy and completes the assembly of the second protomer (Fig 3A).

**Figure 2.**
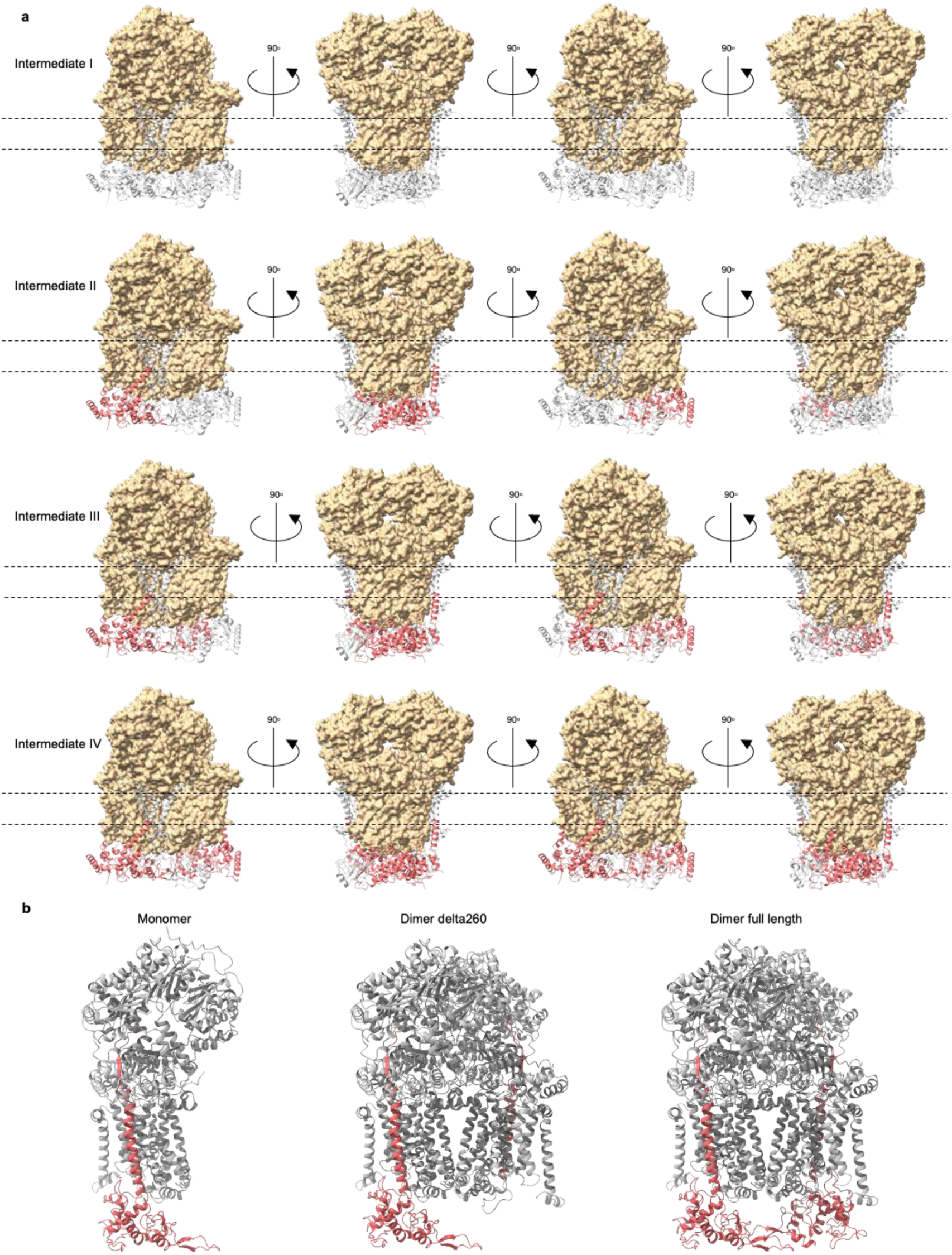
Assembly intermediates of CIII_2_ intermembrane space domain. **a.** Folding steps of the intermembrane space (IMS) domain of CIII_2_ as identified by cryo-EM structures. The core subunits present in all intermediates are shown as surface and coloured in tan, the missing subunits are shown as secondary structures and coloured in grey and the newly folded subunits are shown as secondary structures and coloured in red. The dotted lines highlight the position of the membrane. **b.** AlphaFold3 predictions of Cyt1 folding: Cyt1 is in red and the other subunits are in grey. On the left, a monomer featuring Cor1-2, Cob, Cyt1, Qcr7-8 is shown, in the middle a dimer featuring the same subunits, but where one copy of Cyt1 is truncated in the soluble domain (until residue 260) and on the right a dimer featuring the same subunits, where both copies of Cyt1 are full-length.

**Figure 3.**
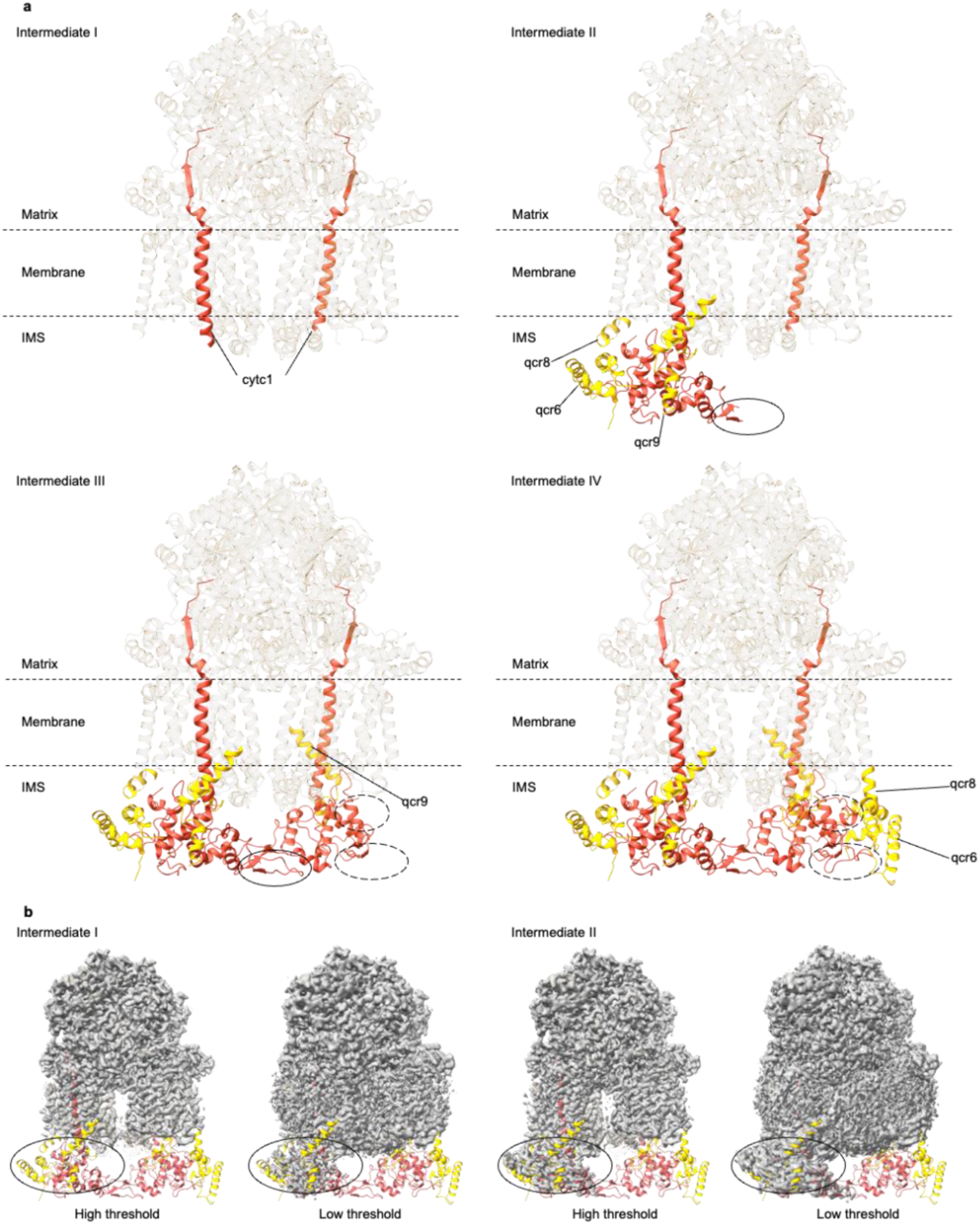
The CIII_2_ intermembrane space domain folds around Cyt1. **a.** The four intermediates of CIII_2_ IMS folding are shown as secondary structures. The core subunits present in all intermediates are partially transparent, coloured in tan. Cyt1 is coloured in red, the other subunits that assemble around it are coloured in yellow and identified by their names. The loops of Cyt1 that fold at a later step compared to the rest of the subunit are highlighted by circles. The 135-146 loop of protomer 1, unfolded in intermediate II and folded in intermediate III, is in the full line circles. The N-terminus until residue 75 and the 198-212 loop of protomer 2, unfolded in intermediate III and folded in intermediate IV, are in the dotted line circles. **b.** Map comparison at high and low thresholds of intermediate I (left) and intermediate II (right). The circle highlights the density in the region of Cyt1 soluble domain. The model of Intermediate IV is fitted to the maps and coloured like the rest of the figure.

The structures suggest that the transmembrane domain of Cyt1 is folded upon assembly in the complex, likely playing a role in the stability of the Intermediate I due to its interactions with both the membrane-embedded subunits and the soluble subunits on the matrix side. Conversely, the soluble domain remains more flexible until it is stabilized by interacting with either Qcr6, as shown by Intermediate II, or with the soluble domain of Cyt1 form the other protomer, as shown by Intermediate III. Although we do not observe any ordered density for the soluble domain of Cyt1 in either protomer of Intermediate I, the flexible soluble domain of Cyt1 is visible at lower map threshold in one protomer of Intermediate I (Fig 3B), confirming that the domain is present in the structure, but it is too flexible to be resolved. Since the assembly of the IMS domain of complex III appears to proceed protomer by protomer, it is likely that the soluble domain of Cyt1 in the second protomer of Intermediate I is more flexible than that of the first protomer, leading to particle alignment on the better resolved protomer and complete loss of density on the other protomer.

When the subunits corresponding to Intermediate I (Cor1-2, Cob, Cyt1, Qcr7-8) are put into AlphaFold3^35^ to predict the structure of the assembly intermediate, the soluble domain of Cyt1 is always predicted as folded, no matter whether one protomer is given as input (Fig 2A left), a perfect dimer (Fig 2A right) or an imperfect dimer where one copy of Cyt1 is given as full length and the other copy is missing the soluble domain sequence (Fig 2A middle). The monomer (Fig 2A left) and the delta260 structures (Fig 2A middle) both lack any interaction partner that would favour the ordering of the soluble domain of Cyt1, nevertheless the domain is still predicted as rigidly folded. This indicates that cryo-EM is crucial to resolve the assembly mechanism of protein complexes, thanks to its ability to separate heterogeneous populations, not just in terms of composition and ordered conformations, but also in terms of partially disordered species that cannot be otherwise identified.

The structures also clarify the role of Qcr6 in complex III assembly: on the one hand Qcr6 is required to stabilize the soluble domain of Cyt1 (Intermediate II), but on the other hand it can initially bind sub-stoichiometrically, as the folding of one Cyt1 copy can itself act as a seed to stabilize the second copy of Cyt1 (Intermediate III). The second copy of Qcr6 only binds afterwards, completing the stabilization of the peripheral loops of Cyt1. These findings are in line with early biochemical studies on Qcr6 deletion in yeast, indicating that the KO strains cannot grow in non-fermentable media at 37° C due to loss of complex III activity. The deletion strains showed decreased Cob absorption and reduced levels of Cyt1, Rip1, Qcr8 and Qcr9 at 37° C, with Cyt1 and Qcr9 already reduced at 30° C^36^. The phenotype can now be interpreted in light of the structural data: deletion of Qcr6 prevents the proper ordering of the soluble domain of Cyt1, in turn also de-stabilizing the interaction with Qcr9 (as observed in Intermediates II and III). Cyt1 and Qcr9 are therefore the first subunits to be affected by the deletion of Qcr6, explaining the phenotype that is already observed at normal growth temperature. It is likely that the higher temperature exacerbates complex III instability, affecting the rest of the membrane-embedded subunits, particularly Qcr8 and Cob. A subsequent study of double KO, where Qcr6 was deleted alongside either Rip1 or Qcr9, identified a faint 500 kDa band positive for the Cor1 and Cor2 proteins^17^. While Qcr6 deletion is known to lead to the accumulation of intermediate Cyt1 (where the second transit peptide is not cleaved)^17,36,37^, this slows down but does not completely halt Cyt1 maturation and it is therefore likely that the mature Cyt1 is assembled into the nascent CIII_2_, but due to the absence of Qcr6 the intermediate becomes unstable and gets turned over, being present at low levels in the membrane.

Our intermediate structures are also in line with the biochemical data on Cyt1 maturation, which indicate that Cyt1 is hemylated prior to processing to the mature form^37^, observed in the context of complex III. Indeed, in all cases where the soluble domain of Cyt1 could be resolved, heme c density was clearly present, even in the second protomer of intermediate III where the distal region of Cyt1, at the interface with Qcr6, is partially unresolved.

### The folding process of the IMS domain is required for supercomplex formation

The classification of supercomplex particles led to the identification of three intermediates, named intermediates I_SC, II_SC and III_SC (Fig 4A). All three intermediates feature intermediates of CIII_2_ bound to fully assembled CIV. More specifically, Intermediate I_SC corresponds to an Intermediate III of CIII_2_ bound to CIV, while intermediates II_SC and III_SC both feature Intermediate_IV of CIII_2_ bound to CIV. By aligning intermediates I_SC and II_SC on CIII_2_ it becomes apparent that the conformation of CIV in intermediate I_SC clashes with the presence of Qcr6, whereas CIV undergoes a minor conformational change in intermediate II_SC that allows positioning of Qcr6 (Fig 4B). This suggests that intermediate III of CIII_2_, missing one Qcr6 subunit, is the earliest CIII_2_ intermediate that interacts with CIV in a supercomplex. In this case, the conformational change in CIV observed in intermediate II_SC is required to have CIII_2_ progress from intermediate II to intermediate III. Intermediate III_SC instead is compositionally identical to intermediate II_SC, but shows a large conformational change of CIV, which pivots around the Cor1- Cox5a interface (Fig 4C). This interaction interface is already established in intermediate I_SC and serves as a hinge to maintain the CIII_2_ to CIV interaction across the different intermediates, until the relative position of the two complexes approximates the fully assembled supercomplex^20,21,33,38^. Notably, the position of CIV in all intermediates and particularly in intermediate III_SC leaves the surface of CIII_2_, where subunits Qcr10 and Rip1 should bind, open to the membrane (Fig 4A bottom view circles), being compatible with the final assembly step of CIII_2_ without requiring further CIV conformational change.

**Figure 4.**
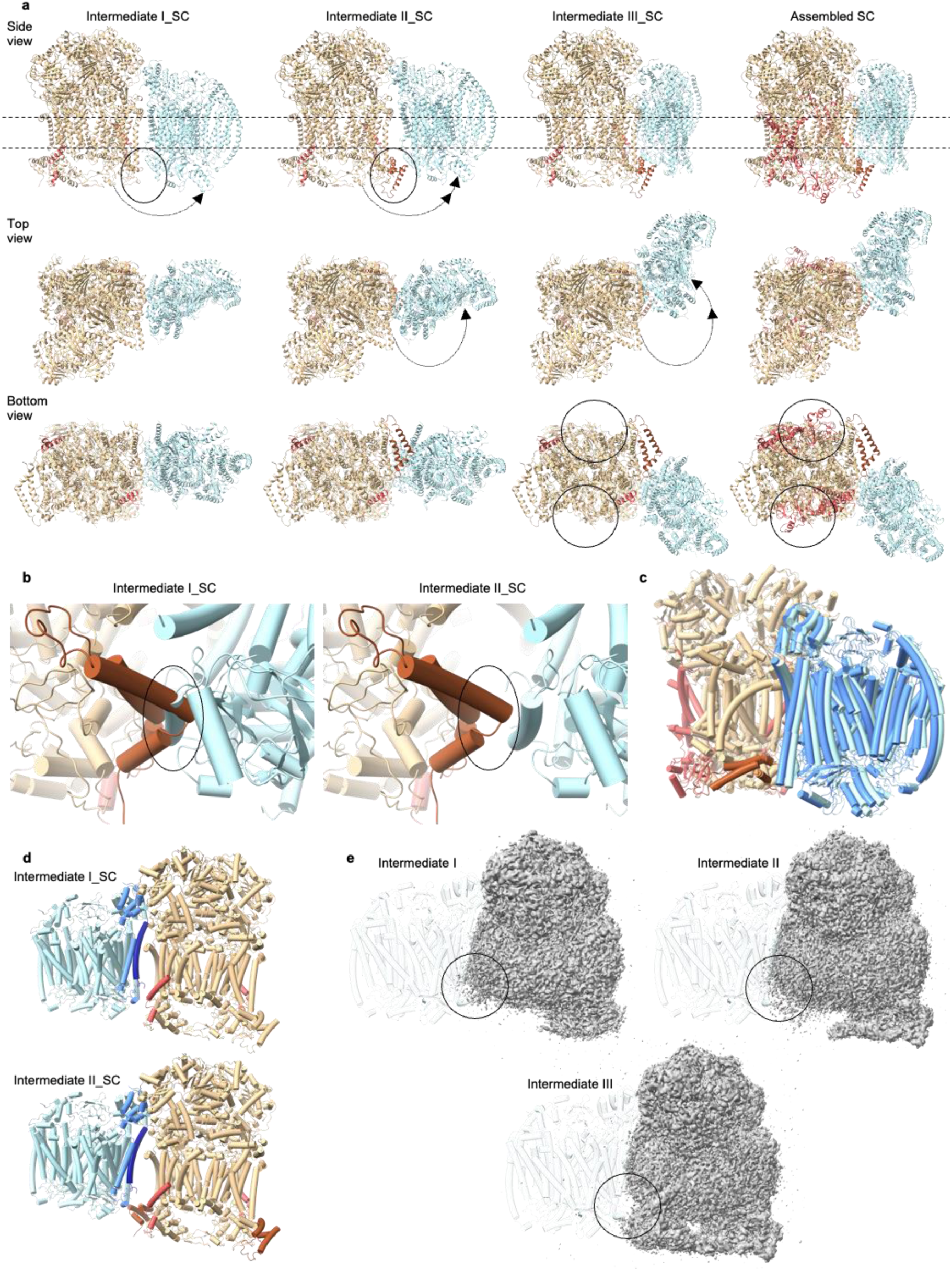
Assembly mechanism of the CIII_2_CIV supercomplex. **a.** The three intermediates of supercomplex (SC) CIII_2_CIV assembly, as identified by the cryo-EM structures, are shown and coloured as in Figure 1, viewed from the side (first line), from the matrix/top (second line) and from the IMS/bottom (third line). The assembled SC corresponds to pdb 9etz. The newly incorporated subunits in each intermediate are coloured in red. The partially folded Qcr9 is shown in red from intermediate I_SC. In the first line, the incorporation of subunit Qcr6 in the protomer at the interface with CIV is highlighted by the circles and the corresponding conformational change of CIV is highlighted by the arrows. Similarly, in the second line the conformational change of CIV is highlighted by the arrows. In the third line, the incorporation of subunits Qcr10 and RIP1 is highlighted by the circles. **b.** The clash between Qcr6 (brown) and Cox5a of CIV (blue) is shown for intermediate I_SC in the circle on the left panel. Due to the conformational change of CIV, the clash is not present in intermediate II_SC on the right panel. **c.** Models of intermediate III_SC and assembled SC (pdb 9etz) are aligned on CIII_2_ and shown as piped helices, similarly to b. CIV of intermediate III_SC is coloured in dark blue, CIV of assembled SC is coloured in ice. **d.** Interacting subunits in intermediates I_SC and II_SC. Cox26 is navy, Cox5a is cornflower blue, Qcr9 is red and Cox6 is brown. **e.** Maps of intermediates I, II and III of CIII2 at low threshold, aligned with CIV from Intermediate I_SC, shown as tube helices and partially transparent. The position of the extra weak density is highlighted by the circle.

In *S cerevisiae* CIII_2_ and CIV can exist as supercomplexes featuring either one or two copies of CIV and their relative proportion is known to depend on the growth conditions, with CIII_2_CIV_2_ being predominant in respiratory conditions and CIII_2_CIV in fermentative conditions^39^: as our cells are respiratory incompetent due to the absence of functional CIII_2_, we expectedly only observe the CIII_2_CIV stoichiometry. Based on the assembly mechanism of CIII_2_, one copy of Qcr6 is needed to stabilize the folding of the first Cyt1 copy and therefore the interaction between CIII_2_ and CIV observed in Intermediate I_SC can only exist in the context of CIII_2_CIV. To attach a second copy of CIV and form CIII_2_CIV_2_, we hypothesize that the interaction observed in Intermediate II_SC could be the entry point for CIV, followed by the same conformational change observed in Intermediate III_SC so that CIV can reach its final position. Notably, we did not observe Intermediates I and II of isolated CIII_2_ in the context of a supercomplex, indicating that the earliest intermediate that forms a supercomplex with CIV already has both copies of Cyt1 fully ordered. As CIII_2_ and CIV interact on the IMS side via Qcr9 in Intermediate I_SC (Fig 4D), it is expected that no supercomplex can form with Intermediate I of CIII_2_. Furthermore, another hypothesis to explain the data is that transiently bound chaperones/assembly factors could prevent CIV attachment to intermediates I and II of CIII_2_. Assembly factors are known to assist the early (Cbp3-4-6, BcaI) and late (BcsI, MzmI) steps of CIII_2_ assembly^9,40–45^ and CCHL, CC_1_HL, Cyc2 and Cyc3 are known to mediate Cyt1 maturation via hemilation^46–50^, while peptidases remove the N-terminal target sequences^37,51,52^. Conversely, there are no known factors that assist Cyt1 incorporation into CIII_2_, nor the assembly of the supernumerary subunits that interact with it on the IMS side. Our maps of intermediates I and II of CIII_2_ show a very weak, diffuse density around the flexible Cyt1, which does not appear in later intermediates. This diffuse density clashes with the position of CIV in all intermediate SC structures, indicating that the putative interaction partner has to be released before CIV can bind (Fig 4E). The density unlikely corresponds to incorrectly classified CIII_2_CIV particles, as no density for the Cox5a soluble domain, interacting with CIII_2_, is observed on the matrix side. This would also be in line with the above-mentioned study of Qcr6 double-KO with Rip1 or Qcr9, featuring a 500 kDa species at low abundance^17^: this intermediate could be initially stabilized by unknown partners, but if these are not eventually replaced by the stable interaction with Qcr6, the intermediate becomes unstable and gets turned over. Further studies will be required to hitherto unknown putative interaction partners, which might be bound either too flexibly or too weakly and/or are bound sub - stoichiometrically, thereby preventing structural elucidation. As the intermediates featuring and missing the diffuse density are separated structurally, but come from the same biochemically heterogeneous preparation, we could not use mass-spectrometry to identify proteins that would be uniquely associated with Intermediates I and II of CIII_2_. Future investigation of assembly intermediates of CIII_2_ will also be required to determine why CIV does not bind to Intermediate II on the side of the ordered protomer.

### Comparison of assembly intermediates of CIII_2_CIV to known supercomplex structures

As shown in Fig 4C, intermediate III_SC closely resembles the position of CIV in the fully assembled CIII_2_CIV, alongside multiple other published structures of CIII_2_CIV and CIII_2_CIV_2_ from *S cerevisiae* in different conditions^20,21,33,38,53,54^ (Fig 5 A-D, CIV in III_SC highlighted by asterisks). The position of CIV in intermediate I_SC (as well as intermediate II_SC) instead resembles the position of CIV from the plant *Vigna radiata* (*V. radiata*)^55^ (Fig 5G). As Cox5a is absent in *V. radiata*, it is possible that after attachment CIV does not change conformation as it cannot pivot around Cox5a, instead remaining in the initial conformation. Interestingly, the position of CIV in intermediates I_SC and III_SC also resemble the position of CIV in the murine supercomplex CIII_2_CIV in the locked (Fig 5H) and unlocked (Fig 5I) conformations, respectively, although the murine CIV is 90° rotated compared to the yeast counterpart^22^ (see Fig 6 asterisk for the 90° rotation, highlighting the position of Cox5a in the yeast and mammalian CIV). The locked CIV conformation corresponds to an assembly intermediate, while the unlocked conformation corresponds to the active form of the supercomplex and the assembly of mammalian CIII_2_CIV depends on the presence of SCAF1, which is not conserved in *S cerevisiae*, as well as in other non-vertebrates^55–58^. The assembly mechanism of murine CIII_2_CIV therefore differs from *S. cerevisiae*. However, the position of CIV relative to CIII_2_ during assembly appears overall conserved. This conservation also appears in CIII_2_CIV from *Toxoplasma gondii* (*T. gondii*)^59^ (Fig 5J). In this organism CIV is roughly twice the size of the opisthokont counterpart and interacts with CIII_2_ via non-conserved subunits, but these interact with Qcr6 in CIII_2_, meaning that the position of CIV around CIII_2_ overall resembles the position of Intermediate II_SC. Since this position leaves the binding sites for Qcr10 and Rip1 open to the membrane in *V. radiata*, *M. musculus* (locked conformation) and *T. gondii*, it is possible that this is the reason for the convergent position of CIV around CIII_2_ during assembly despite the non-conserved rotational orientation of CIV. Notable outliers from this hotspot positions are CIII_2_CIV from *Schizosaccharomyces pombae* (*S. pombae*)^58^ (Fig 5E) and CIII_2_CIV_2_ from *Euglena gracilis* (*E. gracilis*)^57^ (Fig 5F). in the case of *E. gracilis*, the interface between CIII_2_ and CIV is mainly mediated by non-conserved subunits, pointing towards a divergent assembly mechanism that involves these subunits. *S. pombae*, instead, lacks subunit Cox26^58^, involved in the interaction between CIII and CIV in intermediates I_SC and II_SC (Fig 4D). Additionally, the Cor1 residues involved in the interaction with Cox5a in S cerevisiae are not conserved in *S. pombae*^58^. This evolutionary divergence could have resulted in a different assembly mechanism of supercomplex CIII_2_CIV in *S. pombae*, similarly to the case of *E gracilis*, albeit not driven by additional, non-conserved, subunits. Interestingly, it appears that the currently available structures of CIII_2_CIV or CIII_2_CIV_2_ supercomplexes cluster in two groups in terms of CIV position around CIII_2_, one represented by *S. cerevisiae*, mouse (*Mus musculus, M. musculus)* and *V. radiata* and the other represented by *E gracilis* and *S. pombae*.

**Figure 5.**
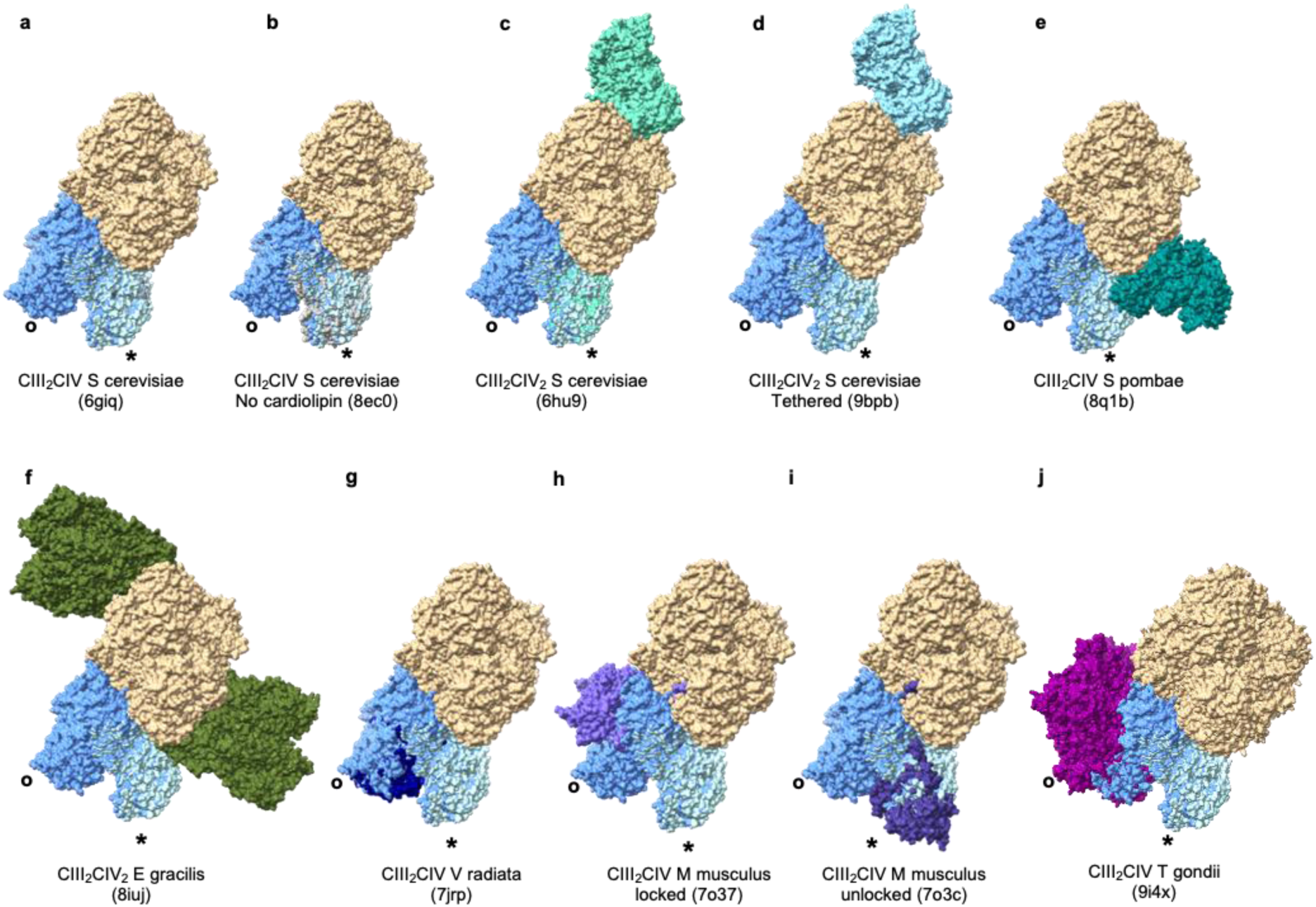
Structural comparison between intermediate SC structures and supercomplex CIII_2_CIV/CIII_2_CIV_2_ from other conditions and species. Intermediates I_SC, III_SC and fully assembled SC (pdb 9etz) are shown from top as surface, aligned on CIII_2_ and coloured in tan (CIII_2_) and dark blue (CIV of intermediates) or ice (CIV of assembled SC). The positions of CIV in the intermediates I_SC and III_SC are further highlighted by circles and asterisks respectively. The models are aligned in each panel on CIII_2_ to different published structures: S cerevisiae CIII_2_CIV (pdb 6giq, CIV grey), S cerevisiae CIII_2_CIV from cardiolipin deficient strain (pdb 8ec0, CIV grey), S cerevisiae CIII_2_CIV_2_ (pdb 6hu9, CIV teal), S cerevisiae CIII_2_CIV_2_ from tethered CIII-CIV (pdb 9bpb, CIV light blue), S pombae CIII_2_CIV_2_ (pdb 8q1b, CIV green), E gracilis CIII_2_CIV_2_ (pdb 8iuj, CIV olive), V radiata CIII_2_CIV (pdb 7jrp, CIV blue), M musculus CIII_2_CIV (pdb 7o37, CIV light purple for locked conformation and pdb 7o3c, CIV dark purple for unlocked conformation), T gondii CIII_2_CIV (pdb 9i4x, CIV dark magenta).

**Figure 6.**
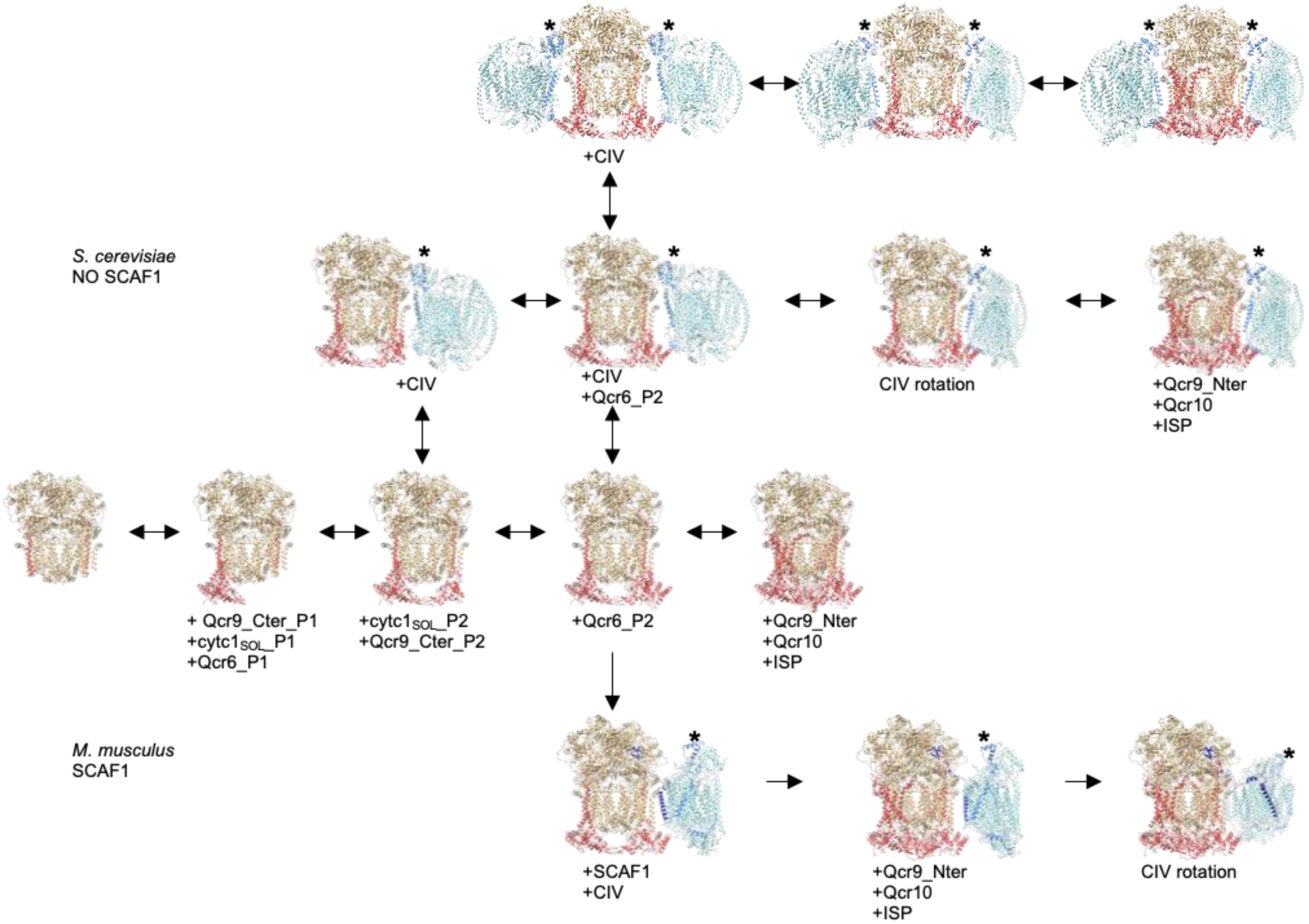
Proposed model of CIII_2_ and CIII_2_CIV assembly mechanism in presence and absence of the assembly factor SCAF1. Assembly mechanism of CIII_2_ IMS domain based on the structures solved in this manuscript. The yeast supercomplex forms from the third intermediate of CIII_2_ IMS folding, while the mammalian supercomplex, forms from the fourth intermediate, as previously described^22^. The added subunits (coloured in red) and conformational changes of CIV are indicated underneath each structure. CIII_2_ and CIV are coloured as in the previous structures. SCAF1 is navy and Cox5a is cornflower blue. The position of Cox5a is further highlighted by the asterisks.

### Cooperative assembly mechanism of CIII_2_ and CIII_2_CIV

Based on our structures, we can now define the intermediate steps of CIII_2_ assembly, focused on the folding of the IMS side of the complex, alongside the cooperative mechanism of SCAF1-independent assembly of CIII_2_CIV starting from an intermediate of CIII_2_ (Fig 6). The assembly of the IMS side of CIII_2_ proceeds asymmetrically, with the addition of one copy of Qcr6 and Qcr9 coupled to the ordering of one Cyt1 soluble domain (Intermediate II). The folding of the first Cyt1 then acts as a scaffold to promote folding of the second Cyt1 soluble domain and the attachment of the second copy of Qcr9 (Intermediate III). This intermediate can then either proceed as an isolated complex or form a supercomplex with CIV (Intermediate I_SC). In either case, the next step consists of the addition of the second copy of Qcr6 (Intermediate IV). Intermediate IV is also observed in a supercomplex with CIV (Intermediate II_SC), which could either be the evolution of Intermediate I_SC or an alternative entry point for supercomplex formation, as well as the potential entry point to add a second copy of CIV to the supercomplex, thereby forming CIII_2_CIV_2_, which we don’t observe in our material. As our structures are solved at steady state, we cannot determine whether intermediates I_SC and II_SC appear consecutively, or represent alternative entry points for supercomplex formation, or if both options occur. Compositionally, Intermediate IV is also present in a different conformation of supercomplex (Intermediate III_SC), where CIV is rotated towards its final position. Given this conformational change, we postulate that Intermediate III_SC follows Intermediate II_SC in the assembly of the supercomplex. Whether CIII_2_ is present in isolation or in supercomplex, the last assembly step is the final maturation of CIII_2_ with the addition of Qcr10, Rip1 and the ordering of Qcr9 N-terminus.

Our structures also corroborate the data on CIII_2_CIV assembly in mouse. Although in mammals CIV binding is mediated by the non-conserved SCAF1 subunit, the assembly mechanism of CIII_2_ itself is conserved and CIII_2_ from the murine assembly intermediate corresponds exactly to Intermediate IV, suggesting that in both species fully assembled CIV forms a supercomplex with late assembly intermediates of CIII_2_. The two pathways eventually diverge due to the presence or absence of SCAF1. In mammals, SCAF1 drives CIV attachment to the CIII_2_ assembly intermediate, and this step is followed by completion of CIII_2_ assembly, before the conformational change of CIV which activates the complex. Conversely, in yeast, where the interface between CIII_2_ and CIV is SCAF1-independent, we first observe the rotation of CIV (Intermediate III_SC) to its final position, prior to completion of CIII_2_ assembly.

As our structures are obtained from steady state and there are no recognizable assembly factors bound to them, we cannot discriminate between *de novo* assembly and turnover. For this reason, Fig. 6 features double arrows. Given the absence of known specific assembly factors, it is possible that assembly and turnover follow the same mechanism, in two opposite directions. Time-resolved studies will be necessary to definitively answer this question. These will require acute re-expression of subunits that were previously KO, to resume de novo assembly, versus acute depletion of subunits from a wild-type background, to accumulate intermediates of disassembly.

CIII_2_ is a highly conserved complex across eukaryotes: the opisthokont (mammalian and yeast) and plant complex features the same protein composition and conformation^20–22,55,58^. The discobal complex (from *E. gracilis*) is also extremely conserved, with the addition of one non-catalytic peripheral subunit^57^. Furthermore, when looking at more available CIII_2_ structures from species that have thus not been presented so far, it is apparent that CIII_2_ is also compositionally and structurally conserved. For example, in the green alga *Chlamydomonas reinhardtii* (*C. reinhardtii*), which is evolutionarily closer to plants as it belongs to the same lineage (Archaeplastida), CIII_2_ is compositionally and structurally identical to opisthokonts^60^. In the SAR lineage (which features Stramenophiles, Alveolata and Rhizaria) instead, the ciliate *Tetrahymena thermophila* (*T. thermophila*) and the apicomplexan *T. gondii* each feature the addition of two short non-catalytic peripheral subunits^59,61^. These findings indicate that the resolved assembly steps, involving conserved subunits across the above-mentioned lineages, are likely conserved in these species, making our model broadly generalizable.

Unlike CIII_2_, CIV appears to display higher compositional diversity, with removal/addition of non-conserved subunits that are often involved in supercomplex formation^57–63^. Although the formation of supercomplex CIII_2_CIV appears in various lineages (Opisthokonta, Archaeplastida, SAR and Discoba), even featuring the emergence of dedicated assembly subunits (like SCAF1 in vertebrates), the conformation of the supercomplex does not appear to be conserved, likely due to CIV variability. This suggests that the assembly steps we observe in *S. cerevisiae* might not be evolutionarily conserved in terms of binding interfaces. However, our structures of the intermediate supercomplexes are also relevant as they provide evidence for the still debated assembly mechanism of respiratory supercomplexes. Two mechanisms have been proposed for supercomplex formation: the cooperative assembly and the plasticity assembly. The cooperative mechanism has been proposed mostly based on biochemical studies in opisthokonts and indicates that intermediates of respiratory complexes can already associate into supercomplexes before their assembly is completed, while in the plasticity model the isolated complexes first fully assemble and then associate into supercomplexes^64^. The structure-based assembly mechanism of the murine supercomplex CIII_2_CIV^22^ already supported the cooperative assembly route and another recently published study also revealed the organization of complex I intermediates into supercomplexes with CIII_2_ and CIV^65^, indicating that the cooperative assembly is not specific for CIII_2_CIV, but can be extended to CI-containing supercomplexes. Our structures perfectly align with the cooperative assembly model, suggesting that this remains a conserved assembly route for CIII_2_CIV in absence of the specific assembly factor SCAF1 and also suggest that for CIII_2_ Intermediate III is the earliest entry route for supercomplex formation. The recent emergence of structural information from evolutionarily divergent species opens a new exciting avenue of research to elucidate the conserved features of supercomplex formation, alongside the confirmation of the cooperative assembly as a universal mechanism of respiratory chain assembly in eukaryotes.

## Materials and Methods

### Yeast strains and growth

Yeast strains had been previously published^10^ and were obtained from the laboratory of Martin Ott. Yeast cultures were grown at 30 °C in YP medium (1% yeast extract and 2% peptone) supplemented with 2% galactose for mitochondrial isolation and subsequent purification.

### Mitochondrial isolation

Cells were lysed with zymolyase (Zymolyase® 20T from Carl ROTH, Art. Nr. 9324.3) treatment followed by osmotic rupture. Mitochondria were then harvested by centrifugation and the total protein content of this fraction was assessed by using the Pierce BCA Protein Assay Kit (Thermo Scientific, ref. 23227).

### Digitonin recrystallization

A modified version of a previously published digitonin recrystallization procedure ^66^ was used to recrystallize digitonin (Carl ROTH, Art. Nr. 4005.4). 25 milliliters of pure ethanol per gram of digitonin were added to the digitonin stock for solubilization. This mixture was solubilized at 75 °C while being stirred under a fume hood. It was then incubated at room temperature for 10 min and then cooled on ice for another 10 min. The material was then centrifuged for 30 min at 7,000g at 4 °C, and the supernatant was discarded. The pellet was redissolved in preheated ethanol (5 ml per initial gram of digitonin) by heating at 75 °C and was vacuum dried for 18 hours. This procedure results in a recovery of around 60% of the initial weight. Digitonin recrystallized in this manner is of sufficient purity to remain stable in purification buffers.

### Purification of CIII_2_ & CIII_2_CIV intermediates

Resuspended mitochondria were solubilized in digitonin and then the material was purified by affinity purification, followed by size exclusion chromatography. Given the presence of multiple heterogenous protein complexes, the final protein concentration was estimated by measuring the 280 nm absorbance signal on a NanoDrop 2000c spectrophotometer (Thermo Scientific). Samples that had a 280 nm absorbance signal between 0.16 and 1.1 were used for grid preparation.

### Grid preparation

Quantifoil® R 2/2 300 copper mesh grids that were manually coated with a continuous carbon layer of approximately 1 nm thickness (https://www.cell.com/iscience/fulltext/S2589-0042(21)00107-3) were glow-discharged for 5 s at 25 mA, 0.39 mbar using the PELCO easiGlow device (Ted Pella, Inc.). Samples were vitrified using a Vitrobot Mark IV (Thermo Fisher). A sample volume of 3 ul was applied to grids after which they were blotted for 5 s using a blot force of 25 at 4 °C, 100% humidity, and were then plunge-frozen in liquid ethane.

### Cryo-EM data acquisition

Micrographs were acquired on a 300kV Titan Krios G4 (Thermo Fisher Scientific) equipped with a Gatan BioContinuum energy filter mounted K3 direct electron detection camera. AFIS (Aberration-Free Image Shift) acquisition was performed using EPU software version 3.8.1.

### Data processing and structure determination

Th cryo-EM micrographs were processed in cryoSPARC^67^. Previously solved structures of S cerevisiae CIII_2_ and CIV were fitted into the Coulomb potential maps and refined in Phenix^68^. The figures were generated in ChimeraX^69^.

## Acknowledgements

We are grateful to Martin Ott for sharing the yeast strains. We gratefully acknowledge the EM training, imaging and access time granted by the life science EM facility of the Ernst-Ruska Center at Forschungszentrum Jülich. EJF is funded by the Emmy Noether Programme (FE 2386/2-1, Deutsche Forschungsgemeinshaft) and the Center for Molecular Medicine Cologne (CMMC, CAP37). We are grateful to Andrea Mattevi, who provided resources to AG through funding by the ERC Advanced Grant MetaQ (grant number 101094471) and Investigator Grant (grant number 28754) by the Associazione Italiana per la Ricerca sul Cancro (AIRC). AG is additionally funded by AIRC through the Postdoctoral Fellowship number 32882 in memory of Angelo and Giancarla Giardina.

